# Cation-chloride cotransporters and the polarity of GABA signaling in mouse hippocampal parvalbumin interneurons

**DOI:** 10.1101/823567

**Authors:** Yo Otsu, Florian Donneger, Eric J Schwartz, Jean Christophe Poncer

**Author notes:** Correspondence should be addressed to Jean Christophe Poncer: INSERM UMR-S 1270, 17 rue du Fer à Moulin, 75005 Paris, France.

## Abstract

Transmembrane chloride gradients govern the efficacy and polarity of GABA signaling in neurons and are usually maintained by the activity of cation chloride cotransporters, such as KCC2 and NKCC1. Whereas their role is well established in cortical principal neurons, it remains poorly documented in GABAergic interneurons. We used complementary electrophysiological approaches to compare the effects of GABAAR activation in adult mouse hippocampal parvalbumin interneurons (PV INs) and pyramidal cells (PCs). Loose cell attached, tight-seal and gramicidin-perforated patch recordings all show GABAAR-mediated transmission is slightly depolarizing and yet inhibitory in both PV INs and PCs. Focal GABA uncaging in whole-cell recordings reveal that KCC2 and NKCC1 are functional in both PV INs and PCs but differentially contribute to transmembrane chloride gradients in their soma and dendrites. Blocking KCC2 function depolarizes the reversal potential of GABAAR-mediated currents in PV INs and PCs, often beyond firing threshold, showing KCC2 is essential to maintain the inhibitory effect of GABAARs. Finally, we show that repetitive 10 Hz activation of GABAARs in both PV INs and PCs leads to a progressive decline of the postsynaptic response independently of the ion flux direction or KCC2 function. This suggests intraneuronal chloride buildup may not predominantly contribute to activity-dependent plasticity of GABAergic synapses in this frequency range. Altogether our data demonstrate similar mechanisms of chloride regulation in mouse hippocampal PV INs and PCs and suggest KCC2 downregulation in the pathology may affect the valence of GABA signaling in both cell types.

**Key point summary:**

- Cation-chloride cotransporters (CCCs) play a critical role in controlling the efficacy and polarity of GABAA receptor (GABAAR)-mediated transmission in the brain, yet their expression and function in GABAergic interneurons has been overlooked.
- We compared the polarity of GABA signaling and the function of CCCs in mouse hippocampal pyramidal neurons and parvalbumin-expressing interneurons.
- Under resting conditions, GABAAR activation was mostly depolarizing and yet inhibitory in both cell types. KCC2 blockade further depolarized the reversal potential of GABAAR-mediated currents often above action potential threshold.
- However, during repetitive GABAAR activation, the postsynaptic response declined independently of the ion flux direction or KCC2 function, suggesting intracellular chloride buildup is not responsible for this form of plasticity.
- Our data demonstrate similar mechanisms of chloride regulation in mouse hippocampal pyramidal neurons and parvalbumin interneurons.

## Introduction

Information representation and processing in the cerebral cortex relies on the dynamic interaction between ensembles of glutamatergic principal neurons and local, highly diversified GABAergic interneurons (Buzsaki, 2010). These interneurons mediate feedforward and/or feedback inhibition onto principal cells (PCs) and thereby control their coordinated activity (Klausberger & Somogyi, 2008). In particular, parvalbumin-expressing interneurons (PV INs), which receive excitatory inputs from both local and distant PCs, in turn provide them with fast perisomatic inhibition (Hu *et al.*, 2014). Fast inhibitory signaling by PV INs controls the timing of principal cell activity (Pouille & Scanziani, 2001) and plays a major role in the generation of rhythmic activities (Klausberger & Somogyi, 2008; Amilhon *et al.*, 2015; Gan *et al.*, 2017) as well as the segregation of PCs into functional assemblies (Agetsuma *et al.*, 2018). However, in addition to excitatory inputs from PCs, PV INs also receive GABAergic innervation from local interneurons (Chamberland & Topolnik, 2012), including some specialized in interneuron inhibition (Gulyas *et al.*, 1996), as well as long-range projecting interneurons (Freund & Antal, 1988). Although GABAergic synapses formed onto PV INs share many properties with those impinging onto principal cells, input-and cell-specific properties were also reported (Chamberland & Topolnik, 2012). For instance, predominant expression of the α1 GABAAR subunit confers PV INs with faster postsynaptic current kinetics as compared to PCs (Gao & Fritschy, 1994; Bartos *et al.*, 2002).

Since GABAARs are predominantly chloride-permeable channels (Bormann *et al.*, 1987), transmembrane chloride gradients also represent a major source of variability for GABA signaling. Cation chloride cotransporters (CCCs) play a critical role in regulating chloride gradients in neurons. Thus, the Na^+^ K^+^ Cl^−^ transporter NKCC1 and the K^+^ Cl^−^ transporter KCC2 are secondary active transporters that regulate intraneuronal chloride using the Na^+^ and K^+^ electrochemical gradients generated by the Na/K-ATPase (Blaesse *et al.*, 2009). Delayed, postnatal KCC2 expression has been shown to contribute to a progressive shift in intraneuronal chloride and the polarity of GABA signaling in cortical PCs *in vitro* (Rivera *et al.*, 1999). *In vivo*, GABA was shown to depolarize immature PCs and yet exert a predominantly inhibitory action on their activity (Kirmse *et al.*, 2015), due to membrane resistance shunting.

However, much less is known regarding chloride handling in GABAergic interneurons. Thus, the reversal potential of GABAAR-mediated currents (E_GABA_) was suggested to be more depolarized in unidentified hippocampal *stratum radiatum* interneurons compared with neighboring PCs (Patenaude *et al.*, 2005). In addition, the driving force of GABAAR-mediated currents was shown to remain unchanged during postnatal maturation, in *stratum oriens* interneurons (Banke & McBain, 2006) but appear to exhibit a hyperpolarizing shift in dentate gyrus basket cells (Sauer & Bartos, 2010). Although most interneurons subtypes were shown to strongly express KCC2 in the adult rat hippocampus (Gulyas *et al.*, 2001), how CCC expression or function control the polarity and efficacy of GABA signaling in these cells remains unknown. One difficulty in addressing this question relates to the diversity and bias of experimental approaches used to evaluate the effect of GABA or chloride transport with minimal perturbation of the neuronal integrity. Here, we used a combination of both invasive and non-invasive *in vitro* electrophysiological approaches to compare GABA signaling in mouse CA1 PV INs and PCs in adult mouse hippocampus. Our results reveal that GABA predominantly exerts depolarizing yet inhibitory actions over both cell types. KCC2 and NKCC1 appear to be functional in both PV INs and PCs even though the two cell types exhibit different somato-dendritic chloride gradients. Finally, we demonstrate that CCCs do not contribute in activity-dependent depression of GABAAR-mediated transmission upon moderate activation frequency (10 Hz). Together our results demonstrate that, in the adult hippocampus, PV INs and PCs both rely on CCC activity to maintain inhibitory GABA signaling.

## Methods

### Animals

*Pvalb*^*tm1(cre)Arbr*^/J mice were crossed with *Gt(ROSA)26Sor*^*tm14(CAG-tdTomato)Hze*^/J (Ai14) reporter mice expressing the red fluorescent protein tdTomato. The genetic background of both *Pvalb*^*tm1(cre)Arbr*^/J and Ai14 mice was C57BL/6J and dual homozygous male or female mice typically aged 35-70 days were used in all experiments. Since we did not observe sex-dependent differences in the biological parameters tested in this study, data from animals of either sex were grouped. All procedures conformed to the International Guidelines on the ethical use of animals, the French Agriculture and Forestry Ministry guidelines for handling animals (decree 87849, licence A 75-05-22) and were approved by the Charles Darwin ethical committee (APAFIS#4018-2015111011588776 v7).

### Immunohistochemistry and imaging

Mice were deeply anesthetized by intraperitoneal injection of ketamine/xylazine (100/20 mg.kg^−1^) and perfused transcardially with oxygenated ice-cold solution containing in mM: 110 choline-Cl, 2.5 KCl, 1.25 NaH_2_PO_4_, 25 NaHCO_3_, 25 glucose, 0.5 CaCl_2_, 7 MgCl_2_, 11.6 ascorbic acid, 3.1 Na pyruvate (~300 mOsm), equilibrated with 95% O_2_-5% CO_2_. Brains were removed and fixed for 4-5 h at 4°C with 4% paraformaldehyde in 0.1M sodium phosphate buffer (pH 7.5) and cryoprotected in 30% sucrose in PBS for an additional 24h. Coronal, 40 μm-thick sections were cut with a cryotome. Free-floating sections were rinsed in PBS and incubated for 4 h in PBS supplemented with 0.5% Triton X-100 and 5% normal goat serum. They were then incubated for 48 h at 4°C with rabbit polyclonal KCC2 antibody (1:400) diluted in PBS supplemented with 0.1% Triton X-100 and 5% normal goat serum before being rinsed in PBS and incubated overnight at 4°C with biotinylated WFA lectin (1:500). The sections were then rinsed in PBS and then incubated for 4h with donkey anti-rabbit Cy5, rinsed in PB and incubated for 40 min with streptavidin Alexa-488. After rinsing in PB, the sections were mounted with Mowiol/Dabco (25 mg.ml^−1^) and stored at 4°C.

KCC2-immunolabeled sections were imaged with a Leica SP5 confocal microscope using a 63× 1.40-N.A. objective with 2X electronic magnification and Ar/Kr laser set at 488, 561 and 633 nm for excitation of Alexa-488, td-tomato and Cy5, respectively. Stacks of 10 optical sections were acquired at a pixel resolution of 0.12 μm and a z-step of 0.29 μm.

### Electrophysiological recordings

Mice were deeply anesthetized by intraperitoneal injection of ketamine/xylazine (100/20 mg.kg^−1^, Sigma-Aldrich) and transcardially perfused with ice-cold solution containing (in mM): 110 choline-Cl, 2.5 KCl, 1.25 NaH_2_PO_4_, 25 NaHCO_3_, 25 glucose, 0.5 CaCl_2_, 7 MgCl_2_, 11.6 ascorbic acid, 3.1 Na pyruvate (~300 mOsm), equilibrated with 95% O_2_-5% CO_2_. Mice were then decapitated and 350 μm-thick parasagittal brain slices were prepared with a vibratome (Microm, Thermo Scientific, France) in the same ice-cold solution and maintained in a humidified interface chamber saturated with 95% O_2_-5% CO_2_ for 10 minutes at 34°C and then at room temperature until use. Artificial cerebrospinal fluid (ACSF) for slice maintenance and recording contained (in mM): 126 NaCl, 26 NaHCO_3_, 10 D-glucose, 3.5 KCl, 1.6 CaCl_2_, 1.2 MgCl_2_, 1.25 NaH_2_PO_4_. For recordings, slices were transferred into a chamber (BadController V; Luigs & Neumann) maintained at 32°C and mounted on an upright microscope (BX51WI; Olympus). Slices were superfused with ACSF at a rate of 2.5 ml.min^−1^.

Loose cell-attached recordings (seal resistance: 15-25 MΩ) were made using 4-6 MΩ borosilicate glass pipettes containing normal ACSF or HEPES-buffered saline containing (in mM): 150 NaCl, 3.5 KCl, 1.6 CaCl_2_, 1.2 MgCl_2_, 10 HEPES, pH 7.4 with NaOH (300 mOsm) in the presence of excitatory transmission blockers (10 μM NBQX and 50 μM D-APV) at a holding potential of 0 mV. Recordings were established by gently pushing the pipette against the membrane of the cell. Signals were filtered at 4 kHz and acquired using pClamp software (Molecular Devices) in voltage clamp mode at a sampling rate of 10-20 kHz.

Tight cell-attached recordings (Perkins, 2006) were performed in the presence of 10 μM NBQX and 50 μM D-APV under current-clamp configuration (I=0 mode) to evaluate the polarity of GABAAR-mediated potentials. Recording pipettes (4-9⍰MΩ) were filled with the HEPES-buffered saline. Seal resistance in the cell-attached mode was >4⍰GΩ. Voltage signals were filtered at 4 kHz and sampled at 10-20 kHz.

For whole-cell recordings, pipettes (3–5 MΩ resistance) were filled with a solution containing (in mM): 115 K-gluconate, 25.4 KCl, 10 HEPES, 10 EGTA, 1.8 MgCl_2_, 2 Mg-ATP, 0.4 Na_3_-GTP, pH 7.4 (290 mOsm) supplemented with Alexa 594 (20 μM) to check cell morphology. Images of the soma and dendrites were acquired at least 15 min after break in, using 535 nm excitation light (CoolLED) to prevent RuBi-GABA uncaging. Cells were voltage-clamped at −60 or −70 mV. Voltage was corrected *post hoc* for liquid junction potential (−11 mV) and voltage drop across the series resistance (<25 MΩ) of the pipette. Currents were filtered at 4 kHz and sampled at 10 kHz.

For gramicidin-perforated patch recordings, the tip of the recording pipette was filled with gramicidin-free solution containing (in mM): 120 KCl, 10 HEPES, 11 EGTA, 1 CaCl_2_, 2 MgCl_2_, 35 KOH, 30 glucose adjusted to pH 7.3 with KOH (300 mOsm). The pipette was then backfilled with the same solution containing 100 μg/ml gramicidin and 20 μM Alexa 488 to verify membrane integrity during the recording. Gramicidin was prepared as a stock solution at 50 mg/ml in DMSO. Pipette resistance was 4-5 MΩ. Cells were voltage-clamped at −70 mV. Recordings were started once series resistance was less than 100 MΩ (52.5±7.6 MΩ for PV INs (n=10) and 69.1±5.8 MΩ for PCs (n=16)). The Donnan potential between the pipette solution and cell cytoplasm was measured (Kim & Trussell, 2007) after spontaneous membrane rupture (11.7±1.1 mV; n = 4). The Donnan potential was partly offset by a liquid junction potential of −4 mV. Therefore, the holding potential in gramicidin-perforated recordings reads 7.7 mV more hyperpolarized than the actual membrane potential. Potentials were corrected for this residual potential and voltage drop across the series resistance of the pipette. Spontaneous action potentials (APs) and resting membrane potential (Vm) were monitored under current clamp configuration (I=0 mode) while E_GABA_ was measured by RuBi-GABA photolysis under voltage clamp. Vm was estimated by averaging membrane potential every 500 ms for 30-60 sec in normal ACSF. Membrane potential in the presence of drugs for photolysis were similarly computed over 30-60 sec (Fig. 6A). The threshold for action potential initiation was determined from the first peak in the third derivative of action potential waveforms averaged from > 4 APs (Henze & Buzsaki, 2001). Currents were filtered at 10 kHz and sampled at 10-20 kHz.

### Photolysis

Photolysis of RuBi-GABA (15 μM) onto parvalbumin positive interneurons (PV INs) or pyramidal cells (PCs) was performed in the presence of 10 μM NBQX, 50 μM D-APV, 2 μM CGP55845 and 1 μM tetrodotoxin (TTX). A 405 nm laser diode beam (Deepstar, Omicron, Photon Lines, France) conducted through a multimode optic fibre and alignment device (Prairie Technologies, Middleton, WI, USA) was set to generate a 3-5 μm spot in the objective focus and directed to the soma or distal dendrites of the recorded neurons. The power of the laser head output was controlled using Omicron Laser Controller v2.97, while trigger and pulse duration were set using pClamp software and a Digidata controller. Photolysis was induced by a 0.5-1 msec pulse at 10 mW on the soma or 3-5 msec at 10 mW on distal dendrites. Series of 15 s voltage steps with a 5 mV increment were applied to the pipette with an inter-episode interval of 40 sec. Laser pulses were delivered at 12 sec after the onset of the voltage step to allow for stabilization of the holding current. The amplitude of GABA-evoked currents was computed as the difference between the current measured over a 4 ms window centered on the peak and the baseline current averaged over 3 ms prior to the laser flash. The distance from soma for dendritic uncaging was measured offline with NeuronJ (Meijering *et al.*, 2004), based on Alexa 594 fluorescence imaging of the recorded neuron.

### Drug application

Isoguvacine (100 μM; Tocris Bioscience) was dissolved in normal ACSF supplemented with 2 μM Alexa 488 to detect regions puffed through a patch pipette using a Picosplitzer III (5 sec at 10 psi). All other drugs were bath applied: NBQX, D-AP5, were from Hello Bio (Bristol, UK). Isoguvacine, RuBi-GABA trimethylphosphine, CGP55845, VU0463271 were from Tocris Bioscience (Bristol, UK). TTX was from Latoxan. All other drugs were from Sigma-Aldrich France. CGP55845, VU0463271 and bumetanide were dissolved in DMSO for stock solutions.

### Data analysis

Electrophysiological data analysis was performed offline using Clampfit 10 (Molecular Devices, USA) and custom routines written in Igor Pro 6 (WaveMetrics, USA).

## Statistical analysis

The results are presented as mean ± SEM throughout the manuscript and in all figures. For statistical analyses, non-parametric Mann-Whitney or Wilcoxon signed-rank tests were used unless Shapiro-Wilk normality test was passed and Student’s t-test could be used. Multiple linear regression analysis was performed using SigmaPlot 12,5 (SPSS). Statistical significance was set at p<0.05.

## Results

### KCC2 expression in hippocampal parvalbumin interneurons

Although the expression and function of CCCs are well characterized in hippocampal principal neurons, whether they are expressed and functional in GABAergic interneurons remains largely unexplored. We used immunohistochemistry in *Pvalb*^*tm1(cre)Arbr/J*^*∷Ai14* mice to investigate KCC2 expression in mouse hippocampal parvalbumin interneurons (PV INs) (Le Roux *et al.*, 2013).

In all hippocampal subfields, KCC2 expression was observed in td-tomato-positive interneurons (Figure 1A). As in PCs, KCC2 immunostaining in PV INs was mostly pericellular, likely reflecting predominant membrane expression (Figure 1B). However, KCC2 expression in PV INs was sometimes difficult to distinguish from that in neighboring PCs. To circumvent this problem, we used extracellular matrix staining to precisely visualize PV IN contours. Hippocampal PV INs somata and proximal dendrites are wrapped by a chondroitin sulfate proteoglycan-rich extracellular matrix, called perineuronal net (PNN) (Hartig *et al.*, 1992). Thus using specific staining of PNNs with Wisteria Floribunda Agglutinin (WFA), KCC2 immunostaining in PV INs could be distinguished from that in adjacent principal neurons as it was surrounded by WFA staining, further confirming KCC2 expression in PV INs (Figure 1B).

These results support the conclusion that KCC2 protein is expressed at the membrane of PV INs in the adult mouse hippocampus.

**Figure 1.**
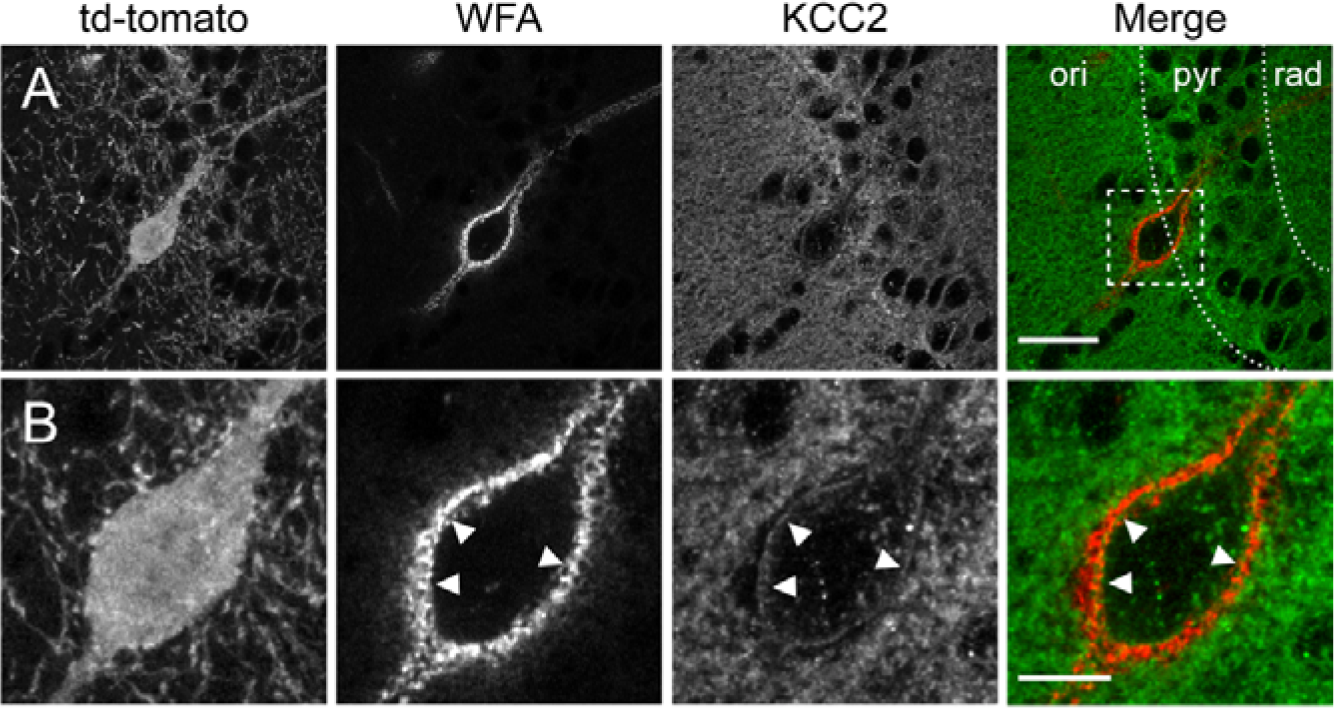
KCC2 labeling of hippocampal CA1 parvalbumin interneurons. A, Representative micrograph (maximum projection of 10 confocal sections over 2.6 μm) of area CA1 of an adult *Pvalb*^*tm1(cre)Arbr/J*^*∷Ai14* mouse hippocampal section immunostained for KCC2 (green) and WFA lectin (red), showing td-tomato expression in a PV IN surrounded by WFA staining on the soma and proximal dendrites. Scale, 30 μm. B, Magnified region boxed in A, showing KCC2 immnostaining in td-tomato expressing PV IN lies just underneath the perineuronal net stained with WFA (arrowheads). Scale, 10 μm.

### Net effect of GABAAR activation on CA1 pyramidal neurons and parvalbumin interneurons

We next asked whether cation-chloride cotransporters are functional in CA1 PV INs and how they influence GABAAR-mediated signaling in these cells. Loose cell-attached recordings allow detection of action potentials from identified neurons with minimal perturbation of their physiology (Llano & Marty, 1995). We first used this approach to evaluate the excitatory vs. inhibitory nature of GABA transmission in CA1 PV INs and neighboring pyramidal neurons (PCs). The effect of GABAAR activation was tested by locally puffing the GABAAR agonist isoguvacin (100 μM, 5 s) onto the soma of the recorded cell. In order to prevent the influence of polysynaptic EPSPs, recordings were performed in the presence of AMPA and NMDA receptor antagonists. Under these conditions however only a few (5 of 27) PV INs exhibited spontaneous firing (Figure 2A). Out of 27 recorded PV INs, isoguvacine induced firing in 1, blocked firing in 5 and had no detectable effect in 21 interneurons. In the presence of the NKCC1 antagonist bumetanide (10 μM), however, isoguvacine suppressed firing in 7 of 14 PV INs, suggesting bumetanide hyperpolarizes E_GABA_ in PV INs. The KCC2 specific antagonist VU0463271 (10 μM), on the other hand, increased the proportion of PV INs that were excited by isoguvacine (3 of 13 cells) while the proportion of cells that were inhibited was similar to that observed in the presence of bumetanide (6 of 13 cells; Figure 2C). These results suggest the effect of GABAA receptor activation is predominantly inhibitory in PV INs and is influenced by the function of both KCC2 and NKCC1.

**Figure 2.**
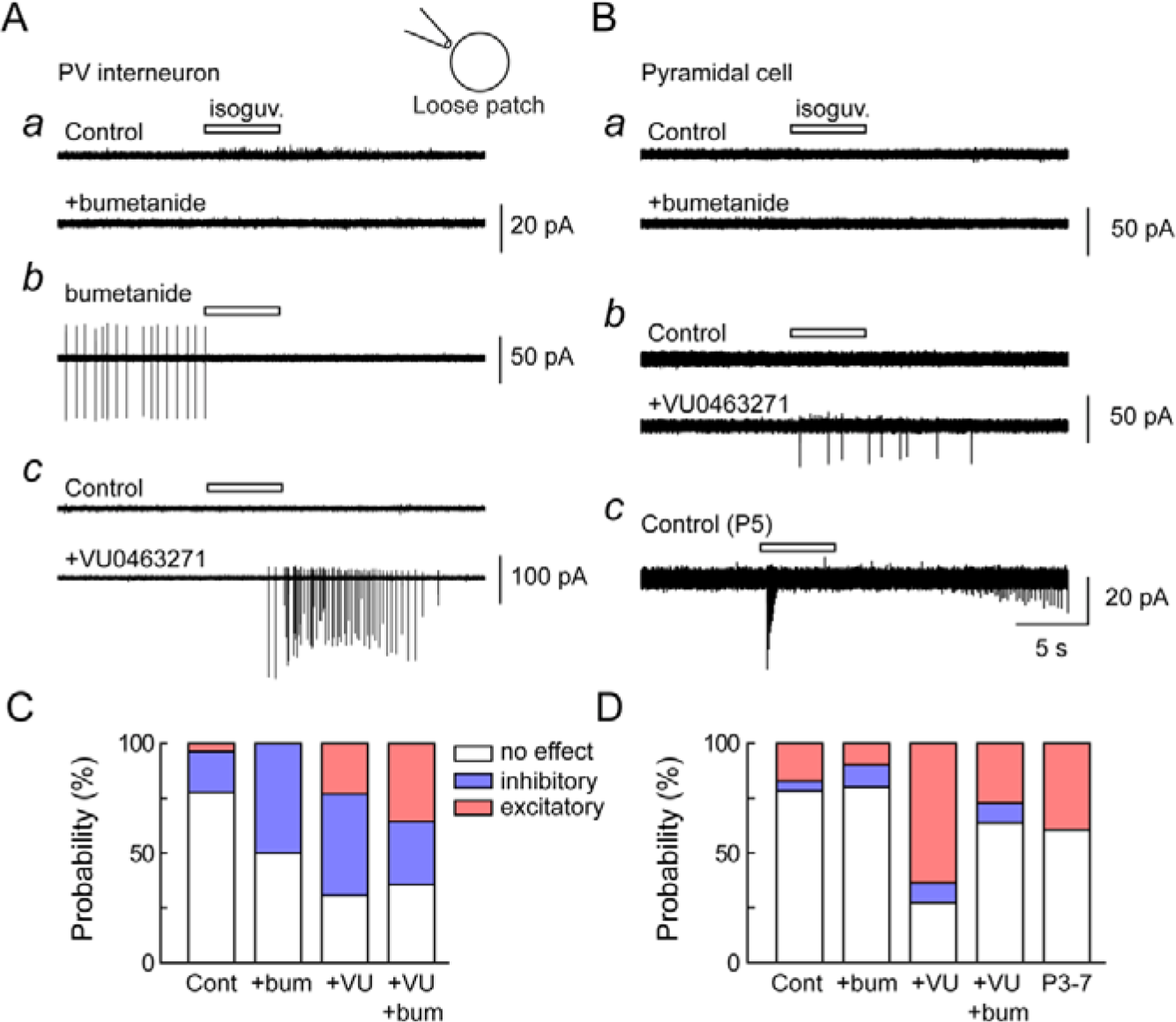
Excitatory and inhibitory actions of GABAA receptor activation in CA1 parvalbumin interneurons and pyramidal cells. A *a*, Representative sections of recordings in loose patch mode of a CA1 PV IN upon brief, focal somatic application of isoguvacine (100 μM, white bar), before and during application of the NKCC1 antagonist bumetanide (10 μM). *b*, same as in *a* in another PV IN during bumetanide application. *c*, same as in *a* and *b*, before and during application of the KCC2 antagonist VU0463271 (10 μM). B *a* and *b*, recordings as in A *a* and *c* from CA1 PCs in P30-P40 mice. *c*, recording showing the effect of somatic isoguvacine application on a CA1 PC from a P5 mouse. C, summary graph showing the proportions of each type response (excitatory, inhibitory or none) recorded upon isoguvacine application in PV INs. D, Same as C for recordings from PCs.

In neighboring PCs, isoguvacine had little effect on firing (5 of 23 cells), mostly owing to the fact that most of them were silent (18 of 23 cells) prior to isoguvacine application, making it difficult to assess the inhibitory or excitatory nature of GABA signaling. Bumetanide had only very little effect on the proportion of pyramidal cells excited (1 of 10 cells) or inhibited (1 of 10 cells) by isoguvacine, whereas VU0463271 induced a large increase in the proportion of excited neurons (7 of 11 cells) (Figure 2D). This excitatory effect was still observed in the presence of both bumetanide and VU0463271 (3 of 11 cells), as with PV INs. In slices from younger (3-7 days old) animals, however, isoguvacine application was sufficient to trigger firing in 26 out of 66 neurons under control conditions, suggesting GABA signaling in PCs was clearly excitatory at this age.

Altogether, these results suggest KCC2 and NKCC1 are functional in both CA1 pyramidal cells and PV INs and influence the efficacy of GABA signaling. However, the high proportion of silent neurons under our recording conditions makes it difficult to draw firm conclusions regarding the polarity of GABA transmission in these cells under physiological conditions.

Tight-seal, cell-attached recordings provide another, minimally invasive approach to detect the polarity of synaptic potentials without rupturing the cell membrane and perturbing transmembrane ionic gradients (Perkins, 2006). In particular, gigaseal recordings allow a fairly reliable measurement of both neuronal resting membrane potential and the polarity (but not the actual amplitude) of synaptic potentials (Mason *et al.*, 2005; Perkins, 2006). We recorded currents evoked by GABAAR activation with isoguvacine in gigaseal mode from both CA1 PV INs and PCs (Figure 3A). In both cell types, isoguvacine-induced potentials were predominantly depolarizing (5 of 7 and 5 of 6 cells, respectively). This proportion was similar to recordings from immature (P3-P7) CA1 pyramidal neurons (6 of 9 cells, Figure 3B). Together, these results suggest that, at least in the absence of glutamatergic drive, both CA1 PCs and PV INs are predominantly depolarized upon GABAAR activation, even though a significant fraction are functionally inhibited, likely due to shunting of their membrane resistance (Staley & Mody, 1992).

**Figure 3.**
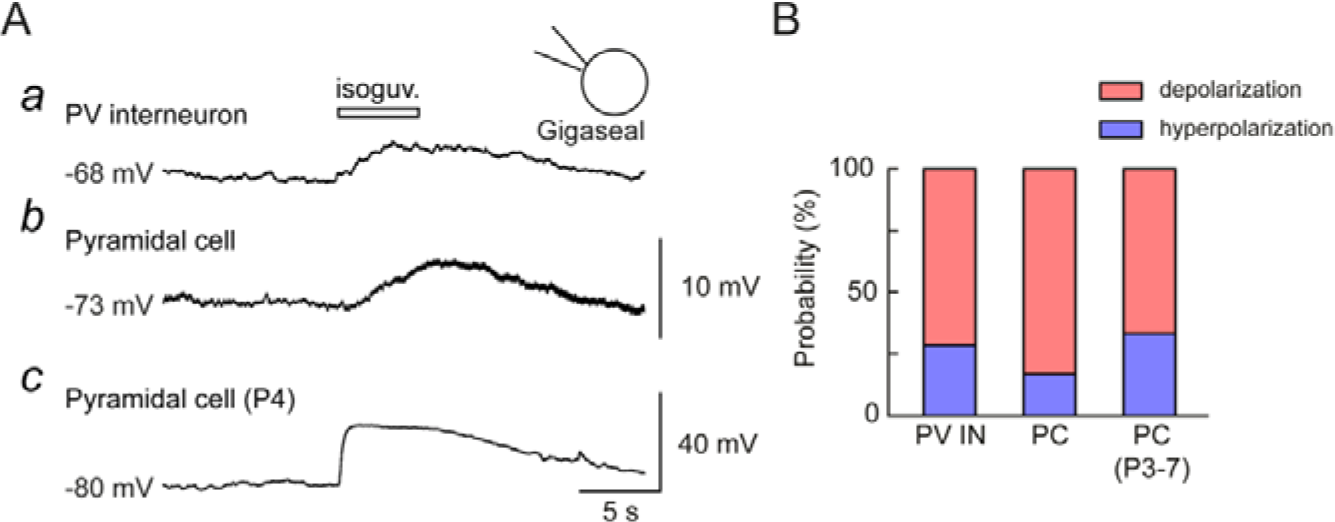
Polarity of GABAAR-mediated potentials in CA1 parvalbumin interneurons and pyramidal cells. A, Representative sections of recordings in gigaseal patch mode of a CA1 PV IN from a P46 mouse (*a*) and a PC from a P33 (*b*) or P4 (*c*) mouse upon brief, focal somatic application of isoguvacine (100 μM, white bar). B, summary graph showing the proportions of each type of response (depolarizing, hyperpolarizing) recorded upon isoguvacine application in each cell type.

### KCC2-mediated chloride extrusion in CA1 parvalbumin interneurons

Transmembrane chloride transport can be directly estimated from whole-cell recordings of GABA-evoked currents while clamping somatic chloride concentration (Khirug *et al.*, 2008; Gauvain *et al.*, 2011). The gradient of the reversal potential of GABAAR-mediated currents (E_GABA_) along the somato-dendritic membrane then reflects actual transmembrane chloride extrusion. We compared E_GABA_ gradients in CA1 PV INs and PCs using local photolysis of RubiGABA (15 μM, 0.5-5 ms laser pulse, see Methods). As in other cortical neurons (Khirug *et al.*, 2008; Gauvain *et al.*, 2011), E_GABA_ in PV INs, was always more depolarized for somatically-evoked currents, as compared to currents evoked onto dendrites 50-250 μm away from the soma (Figure 4). This somato-dendritic gradient (ΔE_GABA_) however was significantly steeper in neighboring PCs as compared with PV INs (−21.1 ± 1.7 *vs* −12.6 ± 0.7 mV/100 μm; 11 dendritic sites in 8 cells and 17 dendritic sites in 9 cells, respectively, p<0.001; Figure 5A-B), suggesting chloride extrusion along dendrites may be less effective in PV INs. However, the effect of KCC2 and NKCC1 blockers on ΔE_GABA_ was not significantly different between the two cell types. Thus, the KCC2 specific antagonist VU046321 produced similar reduction in the somato-dendritic gradient of E_GABA_ in PV INs (−58.5 ± 3.0 %, 15 dendritic sites in 9 cells) and PCs (−57.1 ± 2.7 %, 9 dendritic sites in 7 cells, p=0.770; Figure 4B and 5C). Further application of the NKCC1 antagonist bumetanide also produced similar increase in ΔE_GABA_ in PV INs and PCs (+41.5±7.2% and +45.7±8,5, respectively, as compared to VU046321 only; p=0.67, Figure 5C). This suggests NKCC1 activity may significantly contribute to transmembrane chloride gradients in both cell types, at least upon KCC2 blockade. Altogether, these observations demonstrate chloride extrusion is more efficient along the dendrites of PCs as compared with PV INs and suggest mechanisms other than CCC function may contribute to this difference.

**Figure 4.**
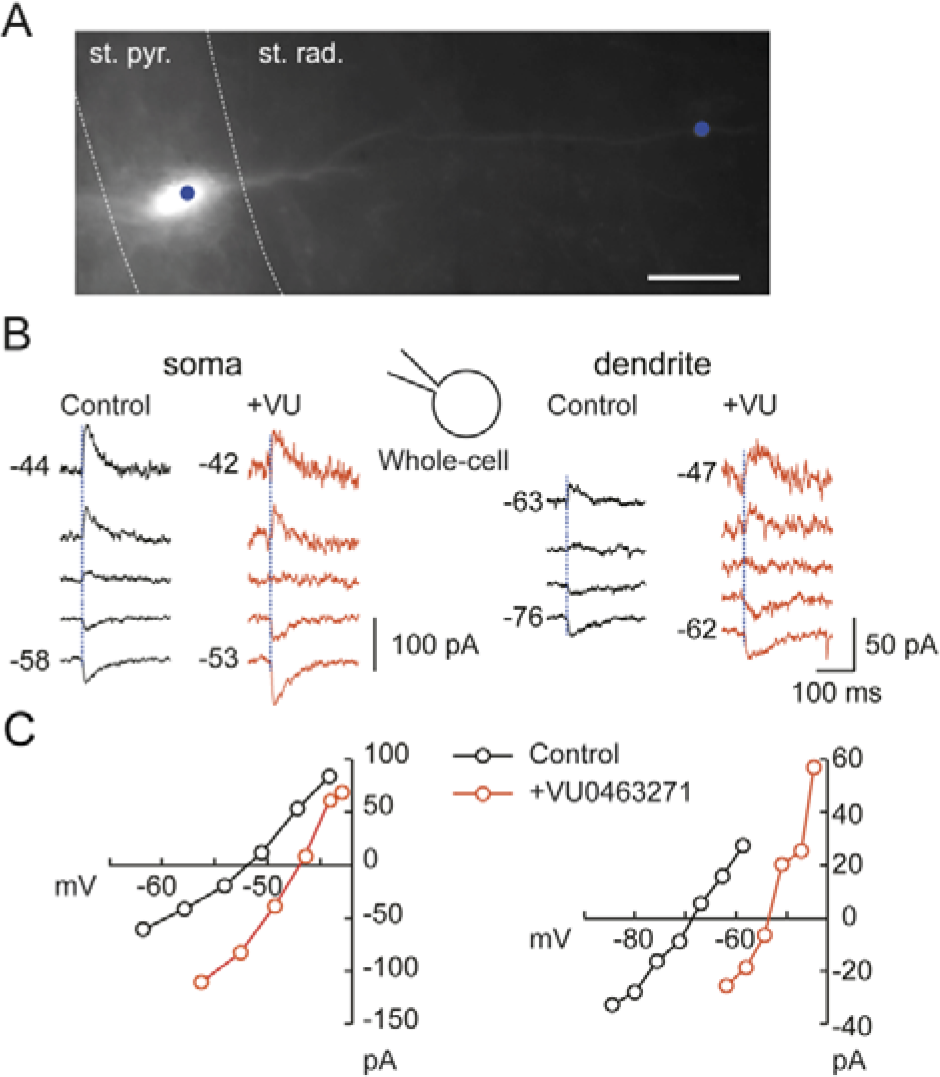
Contribution of KCC2 to transmembrane chloride extrusion in a CA1 parvalbumin interneuron. A, Fluorescence micrograph of a CA1 PV IN from a P37 mouse hippocampal slice, recorded in whole-cell mode and filled with Alex594. The blue spots represent the position and size of the laser beam used for focal RubiGABA photolysis. Scale, 20 μm. B, Currents evoked at varying potentials by focal somatic (left) or dendritic (right) photolysis of RubiGABA in the cell shown in A, before (black) and during (red) application of VU0463271 (10 μM). Numbers of the right represent holding potentials corrected for liquid junction potential and voltage drop across the pipette resistance. C, Current-voltage relations from recordings shown in B showing the different reversal potentials of RubiGABA-evoked currents in the soma vs. dendrites and their depolarizing shift upon KCC2 blockade.

**Figure 5.**
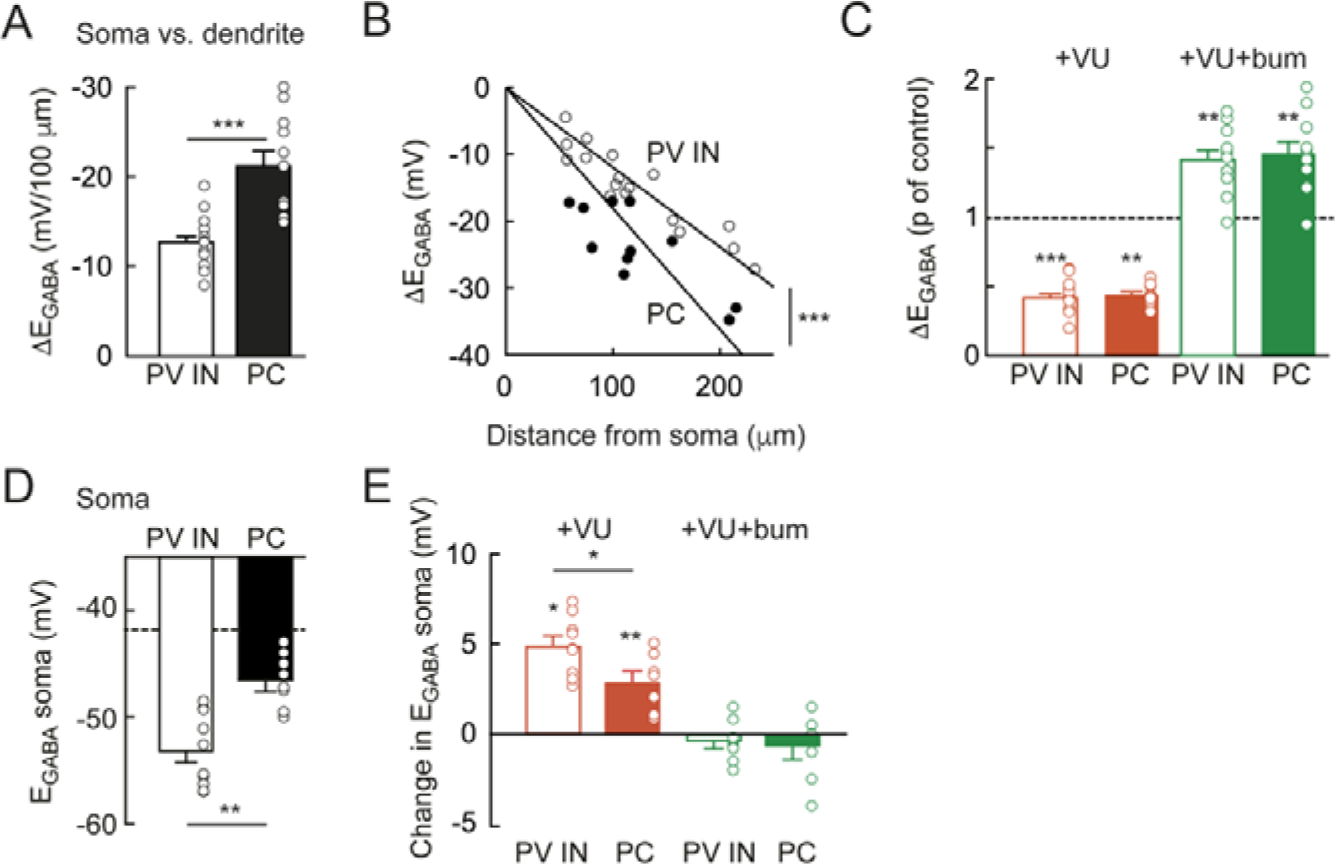
Compared contribution of KCC2 and NKCC1 to somato-dendritic chloride gradients in CA1 parvalbumin interneuron and pyramidal cells. A, Summary graph showing E_GABA_ somatodendritic gradient (ΔE_GABA_) between soma and dendrites normalized by the distance from soma to dendritic photolysis locations. PV IN: n=17 dendritic sites in 9 cells. PC: n=11 dendritic sites in 8 cells. *** Mann Whitney test p<0.001. B, somatodendritic E_GABA_ gradient plotted against the distance from soma to dendritic photolysis locations with superimposed linear regression, showing steeper relation in PCs compared with PV INs. Same data as in A, *** Multiple regression test p<0.001. C, Change in ΔE_GABA_ upon sequential KCC2 (red) and KCC2+NKCC1 blockade (green) by VU0463271 and bumetanide, respectively. The values are normalized to those of the preceding condition (control for VU0463271, VU0463271 for VU0463271+bumetanide). ** and *** Wilcoxon signed-rank test p<0.01 and 0.001, respectively. No significant difference was observed in PC vs PV INs. Same recordings as in A and B. D, Reversal potential (E_GABA_) of currents evoked by somatic RubiGABA uncaging in PV INs and PCs. Same data as in A-C. Dashed line: estimated E_Cl_ based on Nernst equation. ** Mann Whitney test p<0.01. E, Change in somatic E_GABA_upon sequential KCC2 (red) and KCC2+NKCC1 blockade (green) by VU0463271 and bumetanide, respectively. The values are normalized to those of the preceding condition (control for VU0463271, VU0463271 for VU0463271+bumetanide). VU0463271 depolarized E_GABA_ more in PV INs than in PCs. However, further addition of bumetanide had no significant effect. * and ** Wilcoxon rank signed-rank test p<0.05 and 0.01, respectively. * for PV INs vs PCs, Mann Whitney test p<0.05.

Remarkably, although somatic chloride concentration was expected to be clamped by the internal solution of the pipette, somatic E_GABA_ was more hyperpolarized than that estimated by the Nernst equation (−41.3 mV, dashed line in Figure 5D) and more so in PV INs than PCs (53.1±1.1 *vs* 46.5±0.9 mV, n=10 and 8 cells, respectively, p=0. 003; Figure 5D). This suggests that active chloride transport i) may generate transmembrane chloride gradients that do not directly reflect the mean intracellular and extracellular concentrations and ii) is more efficient in the soma of PV INs than in PCs. Consistent with this hypothesis, somatic E_GABA_ was more depolarized upon application of VU0463271 in PV INs than in PCs (+4.9 ± 0.6 *vs* +2.8 ± 0.6 mV, n=9 and 7 cells, respectively, p=0.039; Figure 5E). Further application of bumetanide however had no significant effect on E_GABA_ in either cell type, suggesting NKCC1 does not contribute significantly to somatic transmembrane chloride gradients when intracellular chloride concentration is high (−0.43±0.4 *vs* −0.71±0.7 mV, n=8 and 7 cells, respectively, p= 0.95; Figure 5E). Altogether, our results demonstrate that KCC2 and NKCC1 are functional in PC IN s and contribute to establish steady-state transmembrane chloride gradients. The relative contribution of each transporter to somatic gradients however differ between PV INs and PCs.

In order to assess both E_GABA_ and V_m_ while preventing perturbation of intracellular anion homeostasis, we next used gramicidin-perforated patch recordings. First, we measured V_m_ and tested the effect of pharmacologically blocking the excitatory drive onto CA1 PV INs and PCs, as in experiments shown in Figure 2 and 3. Whereas all PV INs were spontaneously firing at rest (frequency: 4.0±1.4 Hz, n=14 cells), application of AMPA and NMDAR blockers hyperpolarized their membrane potential by 3.5±0.6 mV (Figure 6A and 6Bb). Most PCs (12 out of 15) were also spontaneously firing but had lower firing frequency (0.23±0.1 Hz, p<0.001) and threshold (p=0.006) (Figure 6A and 6Bc). Glutamate receptor antagonists also hyperpolarized CA1 pyramidal cells, yet to a lesser extent than PV INs (0.96±0.3 mV, p=0.001, n= 13 cells of each type, Figure 6Bb), consistent with the latter receiving massive excitatory drive as compared with neighboring pyramidal cells (Gulyas *et al.*, 1999; Takacs *et al.*, 2012). Remarkably, these values of V_m_ measured in the presence of glutamate receptor antagonists were very similar to those derived from gigaseal recordings (71.6±0.6 *vs* −74.9 ± 2.0 mV (n=7) for PV INs and −74.9±1.8 vs. −74.2±0.7 mV (n=8) for PCs; Figure 3). GABAAR - mediated currents were then evoked using focal uncaging of RubiGABA, as above, and E_GABA_ was derived from current-voltage relations of somatically evoked currents (Figure 6C-D). E_GABA_ was significantly more depolarized in PV INs as compared with PCs (−64.1±2.3 *vs* −71.7±0.7 mV, p=0.003, n= 10 and 16 cells, respectively; Figure 6E). However, due to more depolarized V_m_ in PV INs (Figure 6Ba), the driving force of GABAAR-mediated currents at rest was similar in the two cell types (3.8±2.2 *vs* 1.7±1.0 mV, n=10 and 13 cells, respectively; p=0.34; Figure 6F). Also consistent with gigaseal recordings, E_GABA_ was slightly more depolarized than V_m_ both in PV INs and PCs. Application of VU0463271 depolarized somatic E_GABA_ in both cell types (by 12.5±1.6 and 16.1±1.5 mV, in PV INs (n=3) and PCs (n=8), respectively; Figure 6G) whereas further application of bumetanide only slightly hyperpolarized E_GABA_. This suggests that, under steady-state conditions, E_GABA_ in both PV INs and PCs is moderately depolarized as compared with V_m_ and predominantly influenced by the activity of KCC2, whereas NKCC1 contribution is minor. Importantly, however, although GABA may have depolarizing actions in both cells types, its effect is mostly shunting as E_GABA_ is more hyperpolarized than the action potential threshold (Figure 6H). Suppressing KCC2 activity may however be sufficient to depolarize E_GABA_ beyond this threshold, thereby promoting firing, as observed in Figure 2.

**Figure 6.**
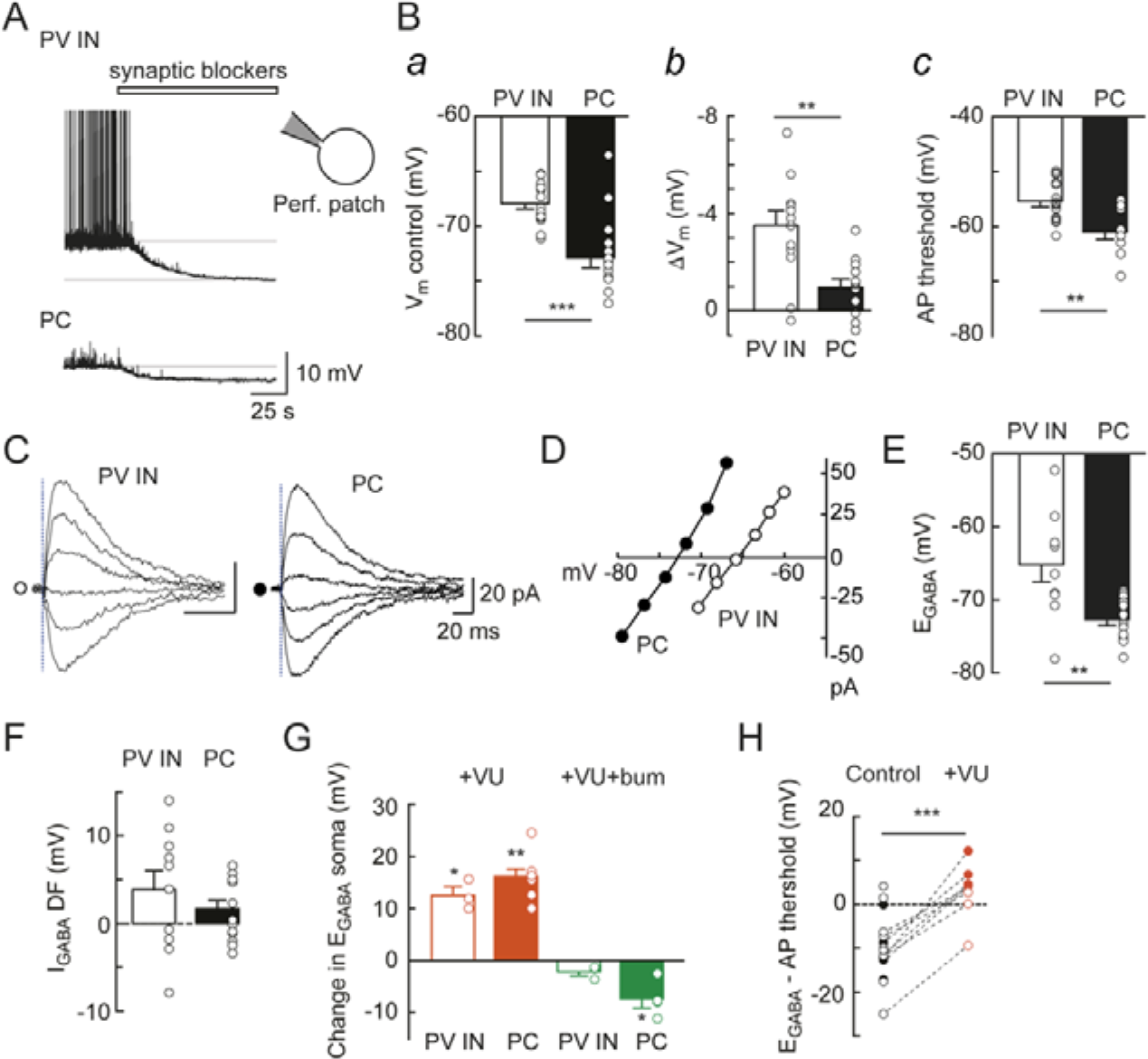
Compared reversal potential and driving force of GABA currents in CA1 parvalbumin interneurons and pyramidal cells. A, Representative current clamp recordings from a CA1 PV IN (top) and PC (bottom) in gramicidin-perforated patch mode, showing the effect of synaptic receptor antagonists (APV, NBQX and CGP55845) and TTX (white bar) on holding potential. Note that the PV IN but not the PC shows spontaneous firing prior to application of the blockers. B, *a*, resting membrane potential measured in 15 CA1 PV INs and 15 PCs prior to application of synaptic blockers. ***, Mann Whitney test p<0.001. *b*, change in membrane potential (ΔV_m_) upon application of synaptic blockers in the same cells as in *a*. **, Mann Whitney test p<0.01. *c*, Action potential threshold measured in spontaneously firing PV INs (n=15) and PCs (n=9). **, Mann Withney test p<0.01. C, Currents evoked at varying potentials by focal somatic photolysis of RubiGABA in a PV IN (left) and a PC (right). The dotted line represents the timing of photolysis. D, corresponding current/voltage relation for the recordings shown in C. Open circles, PV IN. Filled circles, PC. E, Summary graphs showing the reversal potential of somatically evoked GABAAR-mediated currents in 10 CA1 PV INs and 16 PCs. **, Mann Whitney test p<0.01. F, estimated driving force of somatic GABAAR-mediated currents computed by subtracting Vm from E_GABA_, showing GABAARs have slightly depolarizing actions in both PV INs and PCs. G, summary graph showing the change in somatic E_GABA_ upon sequential KCC2 (red) and KCC2+NKCC1 blockade (green) by VU0463271 and bumetanide, respectively. The values are normalized to those of the preceding condition (control for VU0463271, VU0463271 for VU0463271+bumetanide). Addition of bumetanide after VU0463271 had no significant effect on somatic E_GABA_ in either PV INs (n=3) or PCs (n=4). *, paired t-test, p<0.05; **, Wilcoxon signed rank test, p<0.01. H, Difference between E_GABA_ and firing threshold for PV INs (open circles) and PCs (filled circles), before (black) and during (red) application of VU0463271. Dotted lines represent paired data used for statistical comparison (all cells pooled). ***, Paired t-test, p<0.001.

### Dynamic regulation of GABA signaling in CA1 parvalbumin interneurons and pyramidal cells

Repetitive activation of GABAARs has been shown in a variety of neurons to result in activity-dependent depression. This depression was attributed to intracellular chloride buildup (Thompson & Gahwiler, 1989a; Staley & Proctor, 1999; Jedlicka *et al.*, 2011) or to receptor desensitization (Thompson & Gahwiler, 1989c; Jones & Westbrook, 1995) or a combination of both. We compared the contribution of these mechanisms upon repetitive activation of GABAARs in CA1 PV INs and neighboring PCs. In order to exclude presynaptic mechanisms that may contribute to short-term, activity-dependent changes in GABA release (Thompson & Gahwiler, 1989c; Zucker & Regehr, 2002), GABAAR activation was achieved by repetitive (10 Hz), focal uncaging of RubiGABA onto the soma or dendrites of neurons recorded in gramicidin-perforated patch mode (Figure 7A). We then compared the dynamics of GABAAR-mediated currents evoked while holding cells below (−85 to −60 mV) or above (−70 to −50 mV) their reversal potential. Both in PV INs and PCs, the peak amplitude of GABAAR-mediated currents decayed with very similar kinetics during the train, independent of their polarity (soma: τ_inward_=0.11±0.01 vs 0.17±0.04 s^−1^, Mann-Whitney test p=0.291; τ_outward_=0.10±0.02 vs 0.13±0.01 s^−1^, Mann-Whitney test p=0.232; n=6 and 11 cells, respectively) and their site of initiation (soma vs dendrite; 0.242<p<0.695; Figure 7B). This observation suggests the mechanisms involved in the activity-dependent depression of GABAAR-mediated currents during a train of 10 Hz stimulation is unlikely to primarily involve changes in transmembrane ionic gradients. Consistent with this conclusion, application of the KCC2 antagonist VU0463271 had no detectable effect on the decay of outward GABAAR-mediated currents, either in PV INs (τ_outward_ =0.13±0.03 vs 0.11±0.04 s^−1^, paired t-test p=0.363; n=2 cells) or in PCs (τ_outward_=0.11±0.00 vs 0.11±0.01 s^−1^, paired t-test p=0.848; n=4 cells; Figure 7C). We conclude that, at least in our range of current amplitude and stimulation frequency, activity-dependent depression of GABAAR-mediated currents is largely independent of CCC function and does not reflect changes in transmembrane chloride gradients.

**Figure 7.**
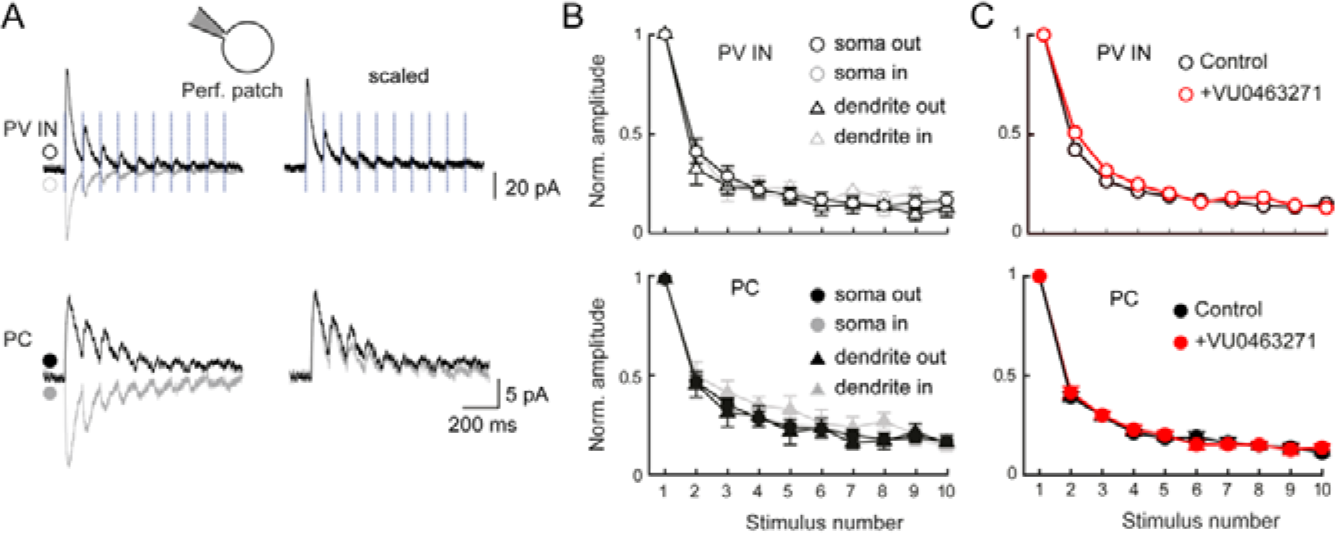
Dynamics of GABAAR-mediated currents in CA1 parvalbumin interneurons and pyramidal cells. A, Representative recordings of currents evoked by 10 Hz somatic photolysis of RubiGABA in a CA1 PV IN and a PC recorded in gramicidin-perforated patch mode and held at potentials above (black) or below (grey) E_GABA_ (PV IN: −60 and −80 mV; PC: −60 and −77 mV). B, Summary graphs showing peak current amplitudes normalized to the peak amplitude of the first current, during a train of RubiGABA photolysis on the soma (circles) or dendrites (triangles) of PV INs (soma: n=6, dendrites: n=3, open symbols) and PCs (soma: n=11, dendrites: n=4, filled symbols). C, Same as in B showing the lack of effect of VU0463271 (10 μM, red symbols) on the decay of the peak amplitude of GABAAR-mediated currents during a 10 Hz somatic RubiGABA photolysis (PV INs, n=2; PCs, n=4).

## Discussion

We have used a combination of approaches to assess and compare the polarity of GABA signaling in adult mouse CA1 hippocampal PCs and PV INs. Our results reveal that the basic mechanisms of steady-state chloride handling controlling GABA transmission are similar in both neuronal types, with a predominantly depolarizing yet inhibitory effect under resting *in vitro* conditions. PV INs and PCs, however, show different behaviors upon intracellular chloride loading, that may reflect differential distribution, regulation or efficacy of cation chloride cotransport along their somato-dendritic axis as well as electrotonic properties. Finally, we have shown that activity-dependent depression of GABAR-mediated transmission is largely independent of the polarity of the evoked currents and the activity of the transporters. This suggests this form of plasticity may not predominantly involve postsynaptic chloride loading, at least under moderate regimes of synaptic activation.

Evaluating the net effect of GABAAR activation in neurons is technically challenging as all experimental approaches may introduce some bias. Classical electrophysiological techniques may induce cell dialysis, compromise membrane integrity or underestimate Donnan potentials between pipette solution and the neuronal cytoplasm (Marty & Neher, 1995). Non-invasive approaches, such as loose-patch recordings, are then often used to assess the polarity and/or the net effect of GABAAR activation on neuronal activity (Deidda *et al.*, 2015; Lozovaya *et al.*, 2019). This approach, however, is only valid when recorded cells display spontaneous firing. As focal application of GABAAR agonists may affect the activity of neighboring neurons and subsequently modify that of the recorded neuron, we performed these recordings in the presence of glutamate receptor antagonists (Fig. 1). In the absence of excitatory drive, however, most recorded neurons (either PV INs or PCs) did not exhibit spontaneous firing and the effect of GABAAR activation could not be tested. These experiments nevertheless showed that GABA agonists mostly inhibit spontaneously firing PV INs. Cell-attached current clamp recordings provide another, minimally invasive approach to evaluate the polarity of synaptic potentials as well as resting membrane potential (Perkins, 2006; Kirmse *et al.*, 2015). Such recordings showed that GABAAR activation mostly induces membrane depolarization in both PV INs and PCs in adult mouse hippocampus, as well as in PCs from immature (P3-7) hippocampus. This observation is supported by gramicidin-perforated patch recordings, which revealed a depolarizing driving force for GABAAR-mediated currents (Fig. 6). Although E_GABA_ was more depolarized in CA1 PV INs than in neighboring PCs, the driving force of GABAAR-mediated currents was remarkably similar, due to a more depolarized resting membrane potential in PV INs. Importantly, under control conditions, such depolarization was not sufficient to reach action potential threshold (Fig. 6H), consistent with a predominantly shunting and inhibitory effect. This observation is in line with earlier studies on cerebellar interneurons (Chavas & Marty, 2003), unidentified hippocampal interneurons (Verheugen *et al.*, 1999; Banke & McBain, 2006) as well as presumptive dentate gyrus PV INs (Sauer & Bartos, 2010).

Very few studies have explored CCC expression in cortical interneurons and data are somewhat controversial, possibly due to the differential expression of distinct isoforms (Uvarov *et al.*, 2007; Markkanen *et al.*, 2014). Thus, KCC2 was shown to be expressed in MGE-derived interneurons earlier than in neighboring pyramidal cells during embryogenesis and to control the termination of their migration (Bortone & Polleux, 2009). However, KCC2 expression and function in specific MGE-derived subtypes in postnatal cortex have not been further explored. In the cerebellum on the contrary, KCC2 expression is very weak in early postnatal presumptive baskets cells and increases postnatally (Simat *et al.*, 2007). In the adult rat hippocampus, KCC2 was shown to be strongly expressed in PV-immunopositive interneurons (Gulyas *et al.*, 2001), consistent with our immunohistochemical data (Fig. 1). Due to the delayed expression of parvalbumin in PV INs (Solbach & Celio, 1991), we could not, however, visualize CA1 PV INs in PVCre∷Ai14 mice in early postnatal mice and therefore could not evaluate the temporal profile of KCC2 expression and function in these cells at earlier stages. In addition, the lack of a specific NKCC1 antibody for immunohistochemistry precluded examination of NKCC1 expression in PV INs and PCs. Our pharmacological data, however, support that both transporters are expressed and functional in both cell types in PV INs and PCs in the adult mouse hippocampus. Thus, the KCC2 antagonist VU0463271 and the NKCC1 antagonist bumetanide had opposing actions on i) the net effect of GABAAR activation on firing (Fig 2), the efficacy of transmembrane chloride export (Fig. 5) and E_GABA_ (Fig. 6) in both PV INs and PCs. Interestingly, however, although KCC2 and NKCC1 blockade had similar effects on somatic E_GABA_ in PV INs and PCs in perforated-patch recordings (Fig. 6G), we observed significant differences when cells were loaded with high intracellular chloride in whole-cell mode. Thus, the somato-dendritic gradient of E_GABA_ was more pronounced in PCs than in PV INs (Fig. 5A-B) and somatic chloride extrusion was more efficient in PV INs than in PCs. These differences are consistent with a differential expression and/or function of KCC2 and NKCC1 along the somato-dendritic axis of the two cell types, with a higher KCC2/NKCC1 function ratio in the soma of PV INs. In addition, differences in the cable properties of PV IN and PC dendrites may also contribute to this difference. Thus, lower membrane resistance of PV IN as compared to PC dendrites (Norenberg *et al.*, 2010)
may induce poorer space clamp of GABAAR-mediated currents evoked onto their distal dendrites. It should also be noted that, whereas the whole-cell evaluation of E_GABA_ gradients is an effective method to assess the efficacy of transmembrane chloride transport (Khirug *et al.*, 2008; Gauvain *et al.*, 2011), it may tend to overestimate steady-state KCC2/NKCC1 function ratio, as it uses high intracellular chloride concentration. This in turn is expected to inhibit the chloride-sensitive with-no-lysine (WNK) STE20 (sterile 20)-like kinases (SPAK) kinases, resulting in reduced NKCC1 and increased KCC2 function (de Los Heros *et al.*, 2014; Friedel *et al.*, 2015; Heubl *et al.*, 2017).

Activity-dependent depression of GABAAR-mediated transmission is well-documented and likely results from a combination of factors involving both pre- and postsynaptic elements (Thompson & Gahwiler, 1989a, b, c). In particular, several studies suggested repetitive activation of GABAARs may lead to postsynaptic chloride loading and a subsequent depolarization of E_GABA_ (Thompson & Gahwiler, 1989a; Kaila *et al.*, 1997; Staley & Proctor, 1999; Magloire *et al.*, 2019). However, these studies used massive chloride loading induced either by multi-quantal IPSCs or prolonged, high-frequency stimulation. Although such intense receptor activation may be relevant to specific, mostly pathological conditions (Magloire *et al.*, 2019), it may not represent the receptor activation at individual, somatic or dendritic sites. Our results from gramidicin-perforated patch recordings instead show that, upon 10 Hz focal Rubi-GABA uncaging for up to 1s, GABAAR-mediated currents decay in amplitude in both PV INs and PCs largely independent of both the direction of the ion flux and KCC2 function (Fig. 7). These results suggest that chloride accumulation during repetitive (10 Hz) activation at single somatic or dendritic sites is not sufficient to significantly affect the driving force of GABAAR-mediated currents, likely owing to the rapid diffusion of chloride ions inside the postsynaptic cytoplasm (Doyon *et al.*, 2011). Instead, since these experiments were performed independent of synaptic stimulation, activity-dependent depression of GABAAR-mediated currents likely reflected receptor desensitization (Jones & Westbrook, 1996; Papke *et al.*, 2011; Gielen *et al.*, 2015). Our results demonstrate this process occurs with a time constant of about 100-130 ms, consistent with the intermediate component of the desensitization kinetics of recombinant α_1_β_1/2_γ_2_L receptors (Papke *et al.*, 2011; Brodzki *et al.*, 2016). Therefore, under physiological regimes of synaptic activity, GABAAR desensitization appears as a major postsynaptic factor acting as a low-pass filter with respect to GABA signaling (Jones & Westbrook, 1996).

CCC expression is altered in a variety of neurological and psychiatric conditions including epilepsy (Palma *et al.*, 2006; Huberfeld *et al.*, 2007; Karlocai *et al.*, 2016; Kourdougli *et al.*, 2017), chronic stress (MacKenzie & Maguire, 2015), Rett syndrome (Duarte *et al.*, 2013; Banerjee *et al.*, 2016; Tang *et al.*, 2016) and autism spectrum disorders (Tyzio *et al.*, 2014). Impaired chloride homeostasis has been suggested to induce paradoxical excitatory GABA signaling and thereby promote anomalous ensemble activities that underlie the pathology. Our data also suggest that KCC2 downregulation may be sufficient to depolarize EGABA above action potential threshold in PV INs (Fig. 6H). In addition, since KCC2 is involved in a variety of molecular interactions with synaptic proteins (Ivakine *et al.*, 2013; Mahadevan *et al.*, 2014), ion channels (Goutierre *et al.*, 2019) and cytoskeleton-related proteins (Li *et al.*, 2007; Gauvain *et al.*, 2011; Chevy *et al.*, 2015; Llano *et al.*, 2015), the loss of its expression also affects several physiological properties beyond the mere control of chloride transport and GABA signaling (Chamma *et al.*, 2012). Thus, KCC2 knockdown in cortical PCs was shown to also profoundly perturb neuronal excitability as well as network activity (Kelley *et al.*, 2018; Goutierre *et al.*, 2019). Since PV INs exert a critical control over the activity of cortical PCs (Pouille & Scanziani, 2001) and shape their rhythmic activities (Klausberger & Somogyi, 2008; Amilhon *et al.*, 2015; Gan *et al.*, 2017), altered CCC expression in these cells would be expected to profoundly perturb cortical rhythmogenesis. As most studies on CCC expression in the pathology lacked cell-subtype resolution, whether and how it is affected in PV INs remains to be fully explored and the consequences on cortical activity should then be further investigated.

## Acknowledgements

We thank X. Marques for expert assistance with confocal imaging, which was performed at the “Photonic Imaging and Image Analysis Platform” of the Institut du Fer à Moulin. We also thank Peter Blaesse for critical reading of the manuscript.

## Author contributions

Y.O. and J.C.P. conceived and designed the research. Y.O. performed all electrophysiological recordings and data analysis with help of E.J.S. who also maintained the mouse colony. F.D. performed immunohistochemistry, confocal imaging and analysis. Y.O and J.C.P. prepared the figures and wrote the paper. All authors approved the final version of the manuscript and agree to be accountable for all aspects of the work in ensuring that questions related to the accuracy or integrity of any part of the work are appropriately investigated and resolved. All persons designated as authors qualify for authorship, and all those who qualify for authorship are listed.

